# Biomarkers to predict steroid resistance in idiopathic nephrotic syndrome: a systematic review

**DOI:** 10.1101/2023.06.21.545865

**Authors:** Carl J May, Nathan P Ford

## Abstract

In this systematic review we have sought to summarise the current knowledge concerning biomarkers that can distinguish between steroid-resistant nephrotic syndrome and steroid-sensitive nephrotic syndrome. Additionally, we aim to select biomarkers that have the best evidence-base and should be prioritised for further research.

Pub med and web of science databases were searched using “steroid resistant nephrotic syndrome AND biomarker”. Papers published between 01/01/2012 and 10/05/2022 were included. Papers that did not compare steroid resistant and steroid sensitive nephrotic syndrome, did not report sensitivity/specificity or area under curve and reviews/letters were excluded. The selected papers were then assessed for bias using the QUADAS-2 tool. The source of the biomarker, cut off, sensitivity/specificity, area under curve and sample size were all extracted. Quality assessment was performed using the BIOCROSS tool.

17 studies were included, comprising 15 case-control studies and 2 cross-sectional studies. Given the rarity of nephrotic syndrome and difficulty in recruiting large cohorts, case-control studies were accepted despite their limitations.

Haptoglobin and suPAR were identified as the most promising biomarkers based on their ability to predict rather than assess steroid resistance in nephrotic syndrome, their respective sample sizes and specificity and sensitivity.

None of the selected papers stated whether the authors were blinded to the patient’s disease when assessing the index test in the cohort.

These candidate biomarkers must now be tested with much larger sample sizes. Using new biobanks such as the one built by the NURTuRE-INS team will be very helpful in this regard.

## Introduction

The kidneys are responsible for many functions vital to sustaining life. They regulate blood pressure, monitor blood pH balance and remove waste products from the blood [1]. The glomerulus is the site of ultrafiltration where small solutes are excreted while proteins and macromolecules are retained [2]. This permselectivity is achieved thanks to the highly specialised structure of the glomerular filtration barrier [3]. The breakdown of the architecture of this barrier leads to runaway proteinuria resulting in the clinical triad of oedema, hypoalbuminemia and proteinuria [4]. This collection of symptoms is termed nephrotic syndrome. Nephrotic syndrome can be genetic or non-genetic. Nephrotic syndrome is the most common glomerular disease of childhood. It has an annual incidence between 1 and 17 cases per 100,000 [5-10]. There are currently over 70 genes that have been implicated in the pathogenesis of nephrotic syndrome. The pathogenesis of non-genetic or idiopathic nephrotic syndrome (INS) is not well understood. The seminal work of Shalhoub *et al and many o*thers since has demonstrated a role for a circulating permeability factor. This factor may be derived from either T-Cells [11, 12], B-Cells [13] or immature myeloid cells [14]. INS is treated with steroids. Steroid sensitive nephrotic syndrome (SSNS) has a very good prognosis with less than 5% progressing to chronic kidney disease [15]. However, between 10 and 20% of patients are steroid resistant (steroid resistant nephrotic syndrome, SRNS) and have a 50% risk of developing end-stage renal failure within 5 years of diagnosis [16]. Even amongst patients who do respond to steroid treatment a subset of these progress to steroid-resistance end-stage renal failure patients require dialysis and or transplant.

It is vital to preserve kidney function by using effective treatments as soon as possible. Currently steroid-resistant patients are identified by their lack of response to a course of steroid treatment. This exposes patients to the unnecessary side-effects of a futile treatment. There is a clear need to be able to differentiate between steroid-sensitive and steroid-resistant patients quickly and accurately.

The use of high-quality biomarkers that can distinguish between steroid-sensitive and steroid-resistant forms of idiopathic nephrotic syndrome would be a paradigm shift for nephrologists and their patients. Instead of being exposed to ultimately useless steroid treatment and enduring the side-effects, steroid-resistant patients could have a simple blood or urine test and proceed to treatment with calcineurin inhibitors.

To summarise what is currently known about potential biomarkers we carried out a systematic literature review. Then applied a quality appraisal tool to identify the most promising biomarker(s) for future more intensive efforts.

## Methods

### Eligibility Criteria

Original research articles that compared biomarkers between known steroid sensitive and steroid resistant nephrotic syndrome patients were included. Review articles, conference proceedings, abstracts and letters to the editor were reviewed as a source for original studies, but excluded from final review. Studies looking at individual candidate biomarkers and panels were included, but studies reliant on kidney biopsies were not included. A decision was made *a priori to limit t*he review to biomarkers from blood, plasma, or urine samples, but not to focus on studies of kidney biopsies, which invasive and painful for the patient. Studies were not excluded based on patient group characteristics beyond having a steroid sensitive and a steroid resistant group.

### Study Screening and Selection

Databases were searched for “Steroid Resistant Nephrotic Syndrome AND Biomarker”. The search results from PubMed and Web of Science were exported to EndNote and screened for duplicates. Studies published between 1^st^ January 2012 until 10^th^ May 2022 were included to focus the review to recent techniques and results. Duplicates were omitted and the titles and abstract of the articles were screened, and the full text of potentially eligible studies was reviewed. Papers were screened by CM without using any automated tools.

### Data Extraction

The sensitivity and specificity or Area Under the Curve (AUC) were extracted for each candidate biomarker or panel. The quality of the data was assessed using BIOCROSS [17] and bias and applicability was scored using QUADAS-2 [18]. CM collected the data and performed the bias and applicability assessment without using any automated tools.

The required characteristics were extracted and tabulated manually by CM and are shown in Figure 2.

### Data Synthesis and Analysis

Data was synthesised and analysed using Excel (Microsoft) and Prism (Graphpad).

### Registration and Protocol

This systematic review was not prospectively registered, and no review protocol has been made available.

## Results

**Figure 1.**
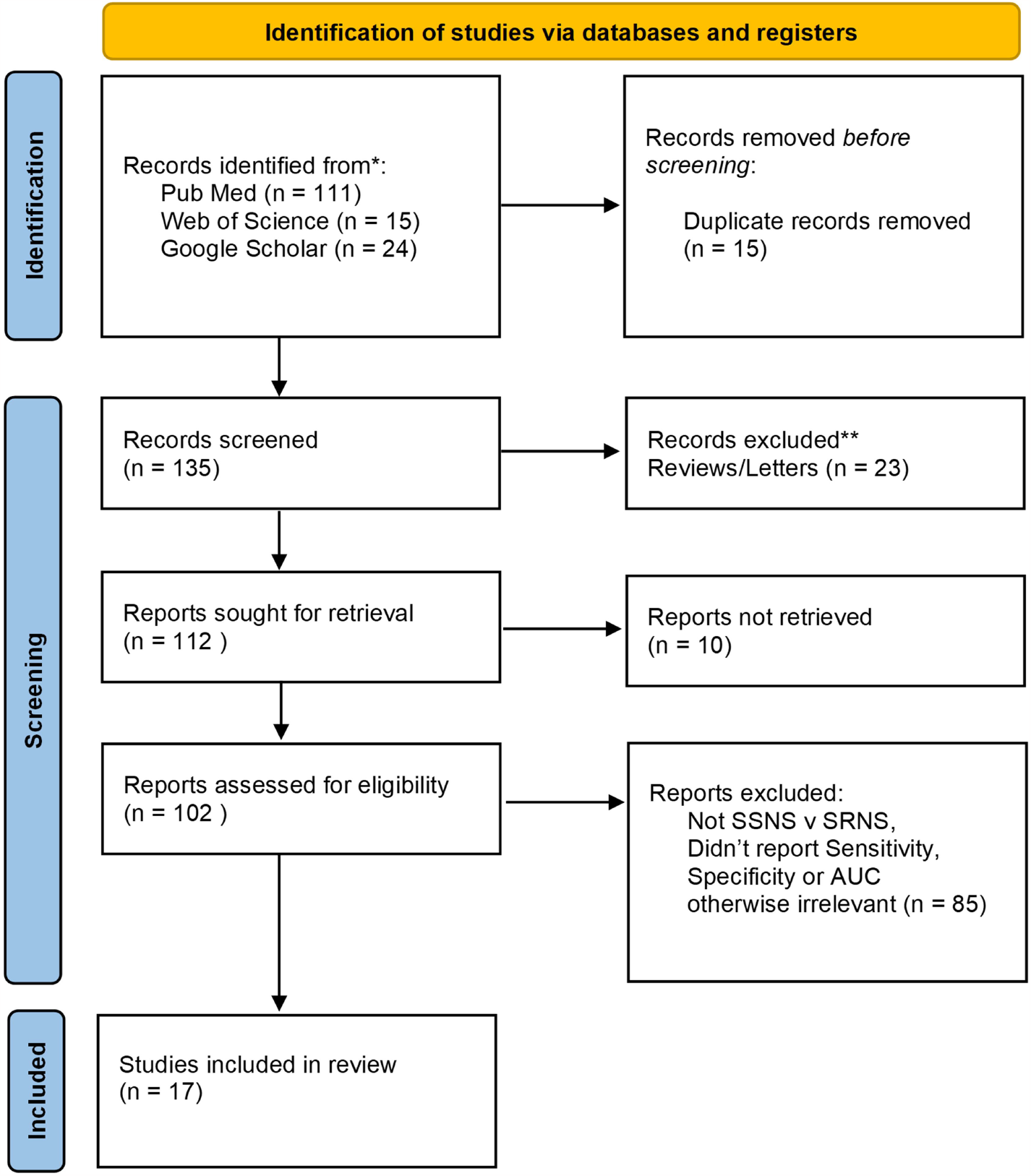
PRISMA Flow Diagram. The PRISMA flow diagram shows the number of studies identified at each stage of the process and why studies were excluded.

Most publications were excluded from the review because they either didn’t compare steroid sensitive patients with steroid resistant patients, or they didn’t report either sensitivity and specificity or AUC. After applying the inclusion/exclusion criteria and screening for eligibility, 17 studies were taken through to review. The most common sample was urine (9 studies) then serum (5 studies) then plasma (2 studies).

**Figure 2.**
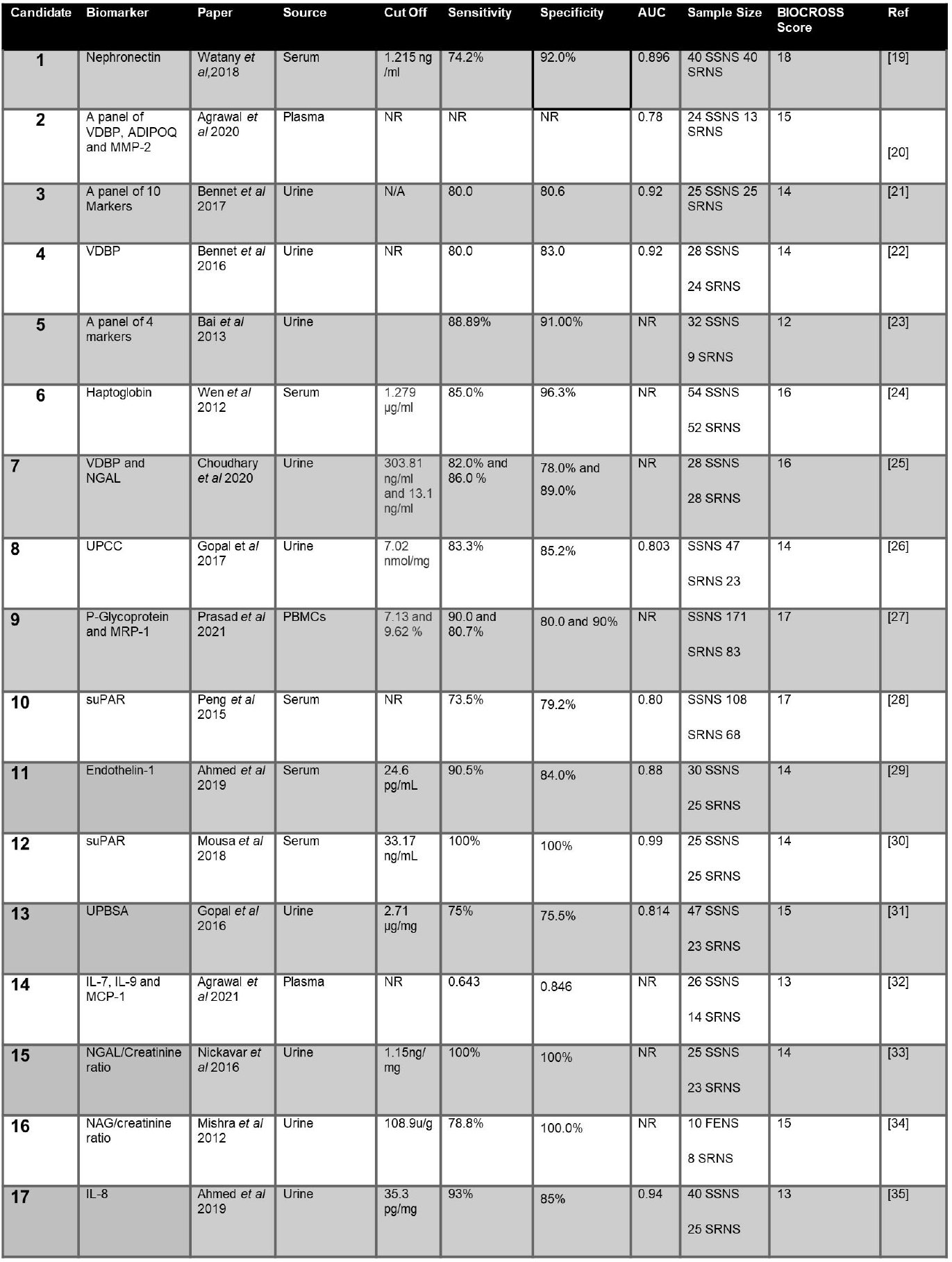
Summary Table of Candidate Biomarkers and Panels. ADIPOQ, adiponectin; MMP-2, matrix metalloproteinase-2. NAG, N-acetyl-beta-D glucosaminodase; NGAL, neutrophil gelatinase-associated lipocalin; SRNS, steroid resistant nephrotic syndrome; SSNS, steroid sensitive nephrotic syndrome; suPAR, soluble urokinase plasminogen activated receptor; UPBSA, urinary protein bound ; UPCC, urinary protein carbonyl content; VDBP, vitamin D binding protein.

The table shown in Figure 1 shows the source paper and key descriptors for the candidate biomarker or panel. The BIOCROSS score indicates the quality of the source article, the higher the number the better the quality [17].

**Figure 3.**
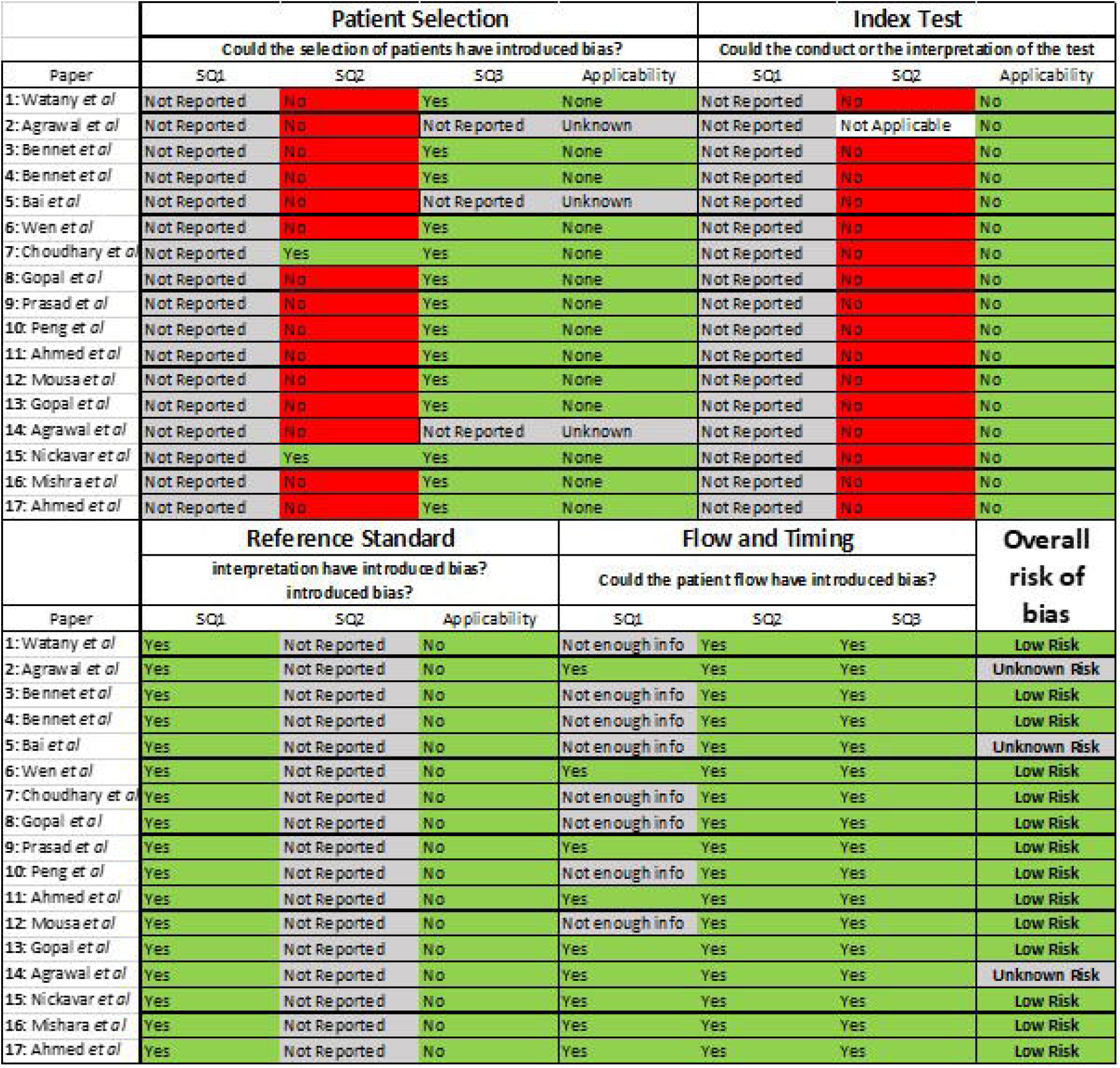
Responses to the QUADAS-2 Assessment. **DOMAIN 1: PATIENT SELECTION Risk of bias: Could the selection of patients have introduced bias?** Signalling question 1: Was a consecutive or random sample of patients enrolled? Signalling question 2: Was a case-control design avoided? Signalling question 3: Did the study avoid inappropriate exclusions? **DOMAIN 2: INDEX TEST Risk of Bias: Could the conduct or interpretation of the index test have introduced bias?** Signalling question 1: Were the index test results interpreted without knowledge of the results of the reference standard? Signalling question 2: If a threshold was used, was it pre-specified? **DOMAIN 3: REFERENCE STANDARD Risk of Bias: Could the reference standard, its conduct, or its interpretation have introduced bias?** Signalling question 1: Is the reference standard likely to correctly classify the target condition? Signalling question 2: Were the reference standard results interpreted without knowledge of the results of the index test? **DOMAIN 4: FLOW AND TIMING Risk of Bias: Could the patient flow have introduced bias? Signalling question 1: Was there an appropriate interval between index test and reference standard? Signalling question 2: Did all patients receive the same reference standard?**

**Figure 4.**
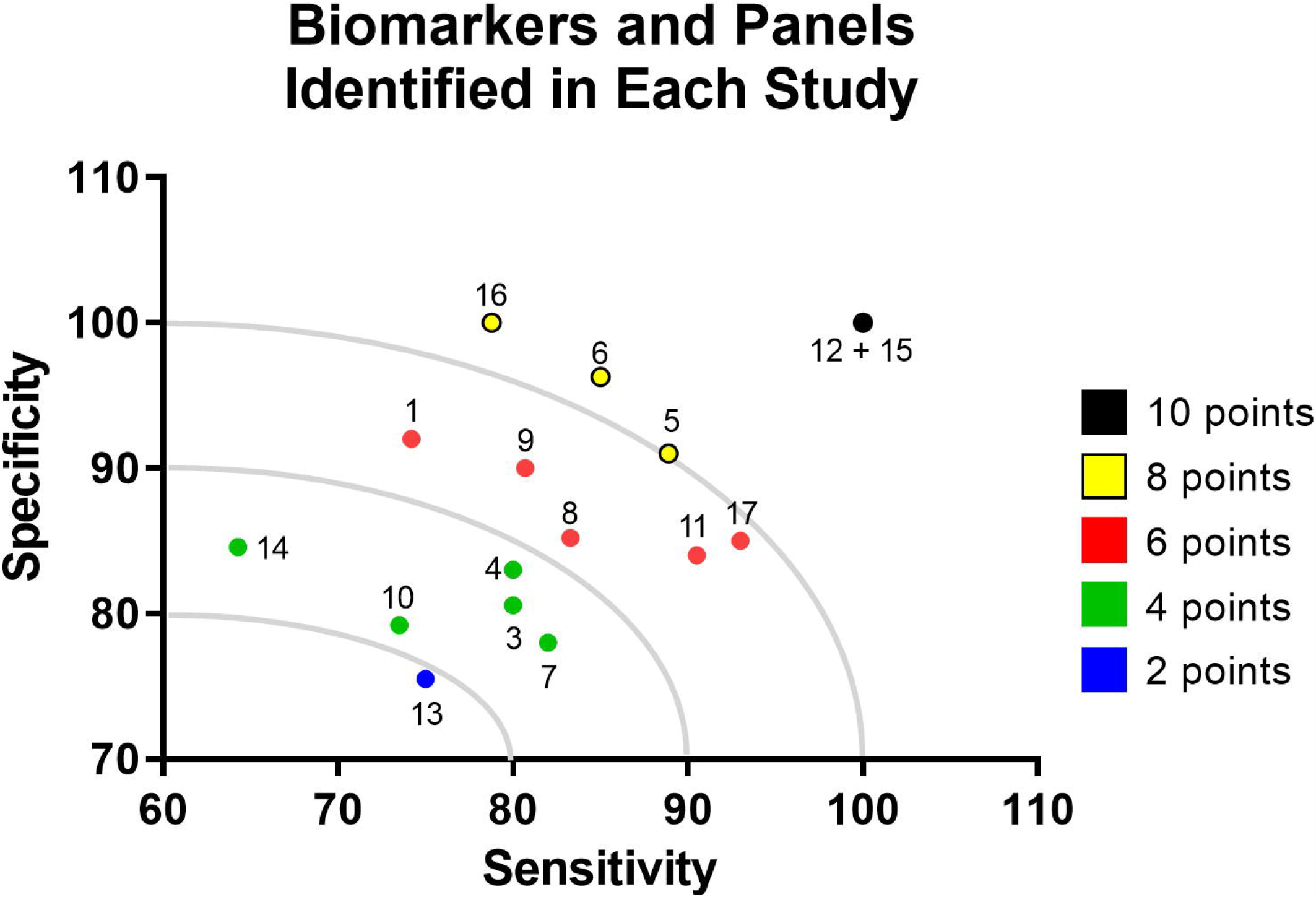
Specificity and Sensitivity of Candidate Biomarkers and Panels. Each biomarker or panel had its specificity plotted against its sensitivity. Points were awarded according to the increasing level of specificity and sensitivity.

### Nephronectin

Nephronectin is a basal lamina protein found in glomerular basement membrane [36]. It is produced by the podocytes and is downregulated following podocyte injury [37]. In glomerular diseases such as focal segmental glomerulosclerosis and membranous nephropathy, nephronectin is also known to be downregulated [37]. Such are these changes in nephronectin expression during and following injury that nephronectin has been identified as a marker of kidney repair following kidney damage [38, 39]. One study, published in 2018, repurposed nephronectin as a marker of kidney repair during the early stages of corticosteroid treatment [19]. For these purposes nephronectin shows promise. However, patients will still be treated with steroids. They may well be removed if there is no evidence of repair (indicated by increased levels of nephronectin); however, ideally a biomarker for steroid resistance would be able to distinguish patients prior to treatment.

### Vitamin D Binding Protein

It has been found that vitamin D deficiency is associated to a greater degree with SRNS compared to SSNS [40]. It has been postulated that this marked vitamin D deficiency is due to the increased urinary loss of VDBP in SRNS versus SSNS [22]. VDBP is sufficiently small to pass through the glomerular filtration barrier. Proximal tubular cells reabsorb the lost VDBP via cubulin and megalin receptors. Hence chronic tubular injury could reduce this reabsorption leading to a greater loss of VDBP in the urine[22]. Hinting at the importance of VDBP as a biomarker of steroid resistance is the presence of VDBP, either on its own or as part of a panel, in four of the seventeen studies.

### ADIPOQ (Adiponectin)

Adiponectin is a hormone, released by adipocytes, that helps to improve insulin sensitivity and is anti-inflammatory [41]. Low levels of adiponectin is correlated with albuminuria in mice and humans [42]. *ADIPOQ knockout m*ice demonstrate significant podocyte injury and albuminuria. Adiponectin therapy in this model restores podocyte foot processes[43]. Elevated levels of serum adiponectin have been reported in patients with FSGS, chronic kidney disease, end-stage renal disease, those on dialysis and transplant recipients [44-46]. Total serum levels of adiponectin rise following the onset of nephrotic syndrome with notable changes in the ratios of the three adiponectin isoforms [47]. A recent study noticed that levels start lower and show a significant decrease following steroid treatment in children with SSNS whereas in children with SRNS levels start higher and increase following treatment [20]. It is under this context that Agrawal proposes using Adiponectin as an early indicator that steroids are working.

### MMP-2

MMP-2 is a metalloproteinase that acts on collagen IV [48]. It is normally expressed by the mesangial cells in the glomerulus; however, during times of inflammation expression levels by the mesangial cells increase and podocytes begin to express MMP-2 [49]. Indeed, increased levels of MMP-2 in the sera have been seen in animal models of chronic kidney disease and in humans with chronic kidney disease [50-52]. MMP-2 can also activate MMP-1 and MMP-9 leading to further extracellular matrix remodelling (ECM) [53]. Increased MMP-2 in the serum and urine has been associated with progressive kidney fibrosis in chronic kidney disease [49, 54-57]. In children with SRNS there is a higher urinary MMP-2/creatinine ratio than in SDNS. This suggests that there may be ECM remodelling in both instances but that in SRNS there is a higher risk of renal fibrosis [58]. One study reported that MMP-2 was elevated in SSNS patients following treatment [20]. Again, this suggests that MMP-2 is useful as an early indicator that steroids may be working but does not help patients avoid steroid exposure altogether.

### NGAL

Neutrophil Gelatinase-Associated Lipocalin (NGAL) is a small 25kDa protein within the lipocalin family [59]. Though initially found in neutrophils, NGAL is expressed by many epithelial cells [60]. It has been widely shown that NGAL expression is upregulated following renal injury and as such is a powerful biomarker for AKI [61-64]. NGAL is a marker for chronic kidney disease progression and is significantly increased in patients with SRNS compared to those with SSNS (AUC0.91 p=<0.0001) [60]. However, it has also been found that calcineurin inhibitors, such as cyclosporine A, can increase NGAL levels [65].

### Fetuin-A

Fetuin-A is a carrier protein that has roles in insulin signalling and protease inhibition [66]. It is central to the pathogenesis of a myriad of conditions including insulin resistance, type 2 diabetes, metabolic disorders, cardiovascular disease and brain disorders [67-70]. In the kidney Fetuin-A protects the integrity of the tissues and levels drop dramatically as chronic kidney disease progresses [71]. Fetuin-A is significantly elevated in the urine during SRNS, suggesting a depletion in the serum leading to a lack of protease inhibition. This is an intriguing hypothesis since there is a body of work supporting the role of a circulating protease in idiopathic nephrotic syndrome [72, 73].

### Prealbumin

Prealbumin can be a sign of a hypercatabolic state often due to increased degradation of muscle mass [74].

### AGP-1

AGP-1 is an acute phase protein released by hepatocytes in response to infection and inflammation [75]. It is generated from active vitamin D [76]. Urinary secretion of AGP-1 in healthy individuals is very low. However urinary secretion is detectable patients in a range of renal diseases including nephrotic syndrome [76].

### Vitamin D Binding Protein

Vitamin D Binding Protein (VDBP) is a circulating protein that binds to vitamin D to create a store of Vitamin D so that rapid vitamin D deficiency can be avoided [77]. In nephrotic syndrome VDBP is lost in the urine along with any vitamin D (or vitamin D metabolites) that is bound to it. This results in low levels of vitamin D as first reported in 1977 [78]. Vitamin D deficiency is more pronounced in SRNS than in SSNS, and VDBP can be used to distinguish between these two conditions [25].

### Haptoglobin

When erythrocytes are lysed the free heme group from haemoglobin can react with molecular oxygen to form superoxide. Haptoglobin binds haemoglobin in the circulation and protects tissues from oxidative damage [79]. Haptoglobin is mainly synthesised in the liver and the lungs, then secreted into the plasma [80]. In addition to its function as an antioxidant, haptoglobin also plays roles in angiogenesis, immunoregulation, the inhibition of nitric oxide and it stimulates tissue repair [81]. There is a significant increase in serum haptoglobin in SRNS patients versus SSNS patients [24].

### Urinary Protein Carbonyl Content UPCC

Chronic oxidative stress can result in systemic inflammation. In turn this can lead to secretion of pro-inflammatory cytokines and an exacerbation of proteinuria [82]. An imbalance between oxidants and anti-oxidants has long been known to exist in idiopathic nephrotic syndrome [83]. Since the oxidative stress is said to be higher in SRNS Vs SSNS then it stands to reason that there would be more UPCC [84], and this has been reported [31]. The pro-inflammatory cytokines induced in response to chronic oxidative stress can damage podocytes [85, 86].

### P-Glycoprotein

P-glycoprotein is encoded by the *MDR-1 gene, it i*s a transporter that causes the efflux of toxins and drugs that are between 300 and 2000 kDa [87]. Cyclosporine is a P-glycoprotein antagonist that reduces the activity of the transporter allowing corticosteroids to accumulate within the cell to achieve a therapeutic effect [88]. The activity of P-glycoprotein was found to be significantly higher in the peripheral blood mononuclear cells (PBMCs) of steroid resistant patients compared to steroid sensitive patients [89]. P-glycoprotein expression is higher on the lymphocytes of steroid resistant patients versus steroid sensitive [90]. Additionally, glomerular expression of P-glycoprotein is significantly increased in those who frequently relapse and or steroid resistant or steroid dependent [91]. Genetic differences affecting the activity of P-glycoprotein can have an impact on the outcome of drug therapy [92] and a significant increase in the prevalence of a SNP in the *MDR1 gene (G267*7T/A) amongst SRNS patients compared to SSNS has been reported [93].

### MRP-1

MRP-1 is pump responsible for the efflux of drugs acting on similar substrates to P-glycoprotein, but with a preference for heavy metal anions and toxins out of cells [94]. In contrast with most ABC transporters that are located on the apical membrane of cells and pump out into the urine or bile, MRP-1 is located on the basolateral membrane and pumps out into the interstitium [95, 96]. It is expressed throughout the body but particularly highly in PBMCs and the kidney [97]. Increased expression of MRP-1 is known to be associated with SRNS and can be assayed for [27].

### suPAR

The soluble urokinase plasminogen activator receptor (suPAR) fulfils criteria of a circulating protein signalling to the kidney.It can readily be found in the circulation. It is generated by immature myeloid cells [14]. It can act directly on the podocyte via interaction with αvβIII integrin [98] or activates proximal tubular cell mitochondria [99]. The former has a deleterious effect on the podocytes, leading to foot process effacement and downregulation of podocin and nephrin [98]. This can affect the structure and function of the glomerulus [100-103]. Urinary suPAR is known to increase in FSGS and positively correlates with disease severity [104, 105]. Additionally it has been shown to be able to predict recurrence of disease in transplanted patients [106]. Urinary and serum levels of suPAR can also stratify cases of minimal change disease and FSGS [107]. There is increasing evidence supporting its utility as a biomarker of steroid resistance in FSGS [108, 109].

### Interleukin-7 (IL-7)

IL-7 is secreted by T cells. T cells were postulated to be the source of the illusive circulating factor driving INS almost fifty years ago [110]. IL-7 is a cytokine that supports host defence by regulating the homeostasis of the cells of the immune system such that congenital deficiency of IL-7 leads to severe immunodeficiency [111]. In the mouse model of Adriamycin nephropathy, IL-7 has been shown to lead to impaired barrier function, podocyte apoptosis, impaired activation of nephrin and actin cytoskeleton dysregulation [112].

### Interleukin-9 (IL-9)

IL-9, also released by T cells, seems to act antagonistically to IL-7 on the podocyte in nephrotic syndrome pathogenesis. IL-9 can dramatically improve glomerular function in the Adriamycin nephropathy model. However, it is known to increase in the serum of patients with primary FSGS hence it’s utility here as a biomarker [113].

### Interleukin-8 (IL-8)

IL-8 is released by T-cells or resident kidney cells in response to pro-inflammatory stimuli [114]. Within the kidney IL-8 is produced by mesangial cells [115], podocytes [116] and tubular epithelial cells [117]. IL-8 is known to affect the functioning of the glomerular basement membrane [118]. It has been shown in rats that IL-8 treatment decreases the synthesis of heparin sulfate proteoglycans leading to proteinuria [119].

**Figure 5.**
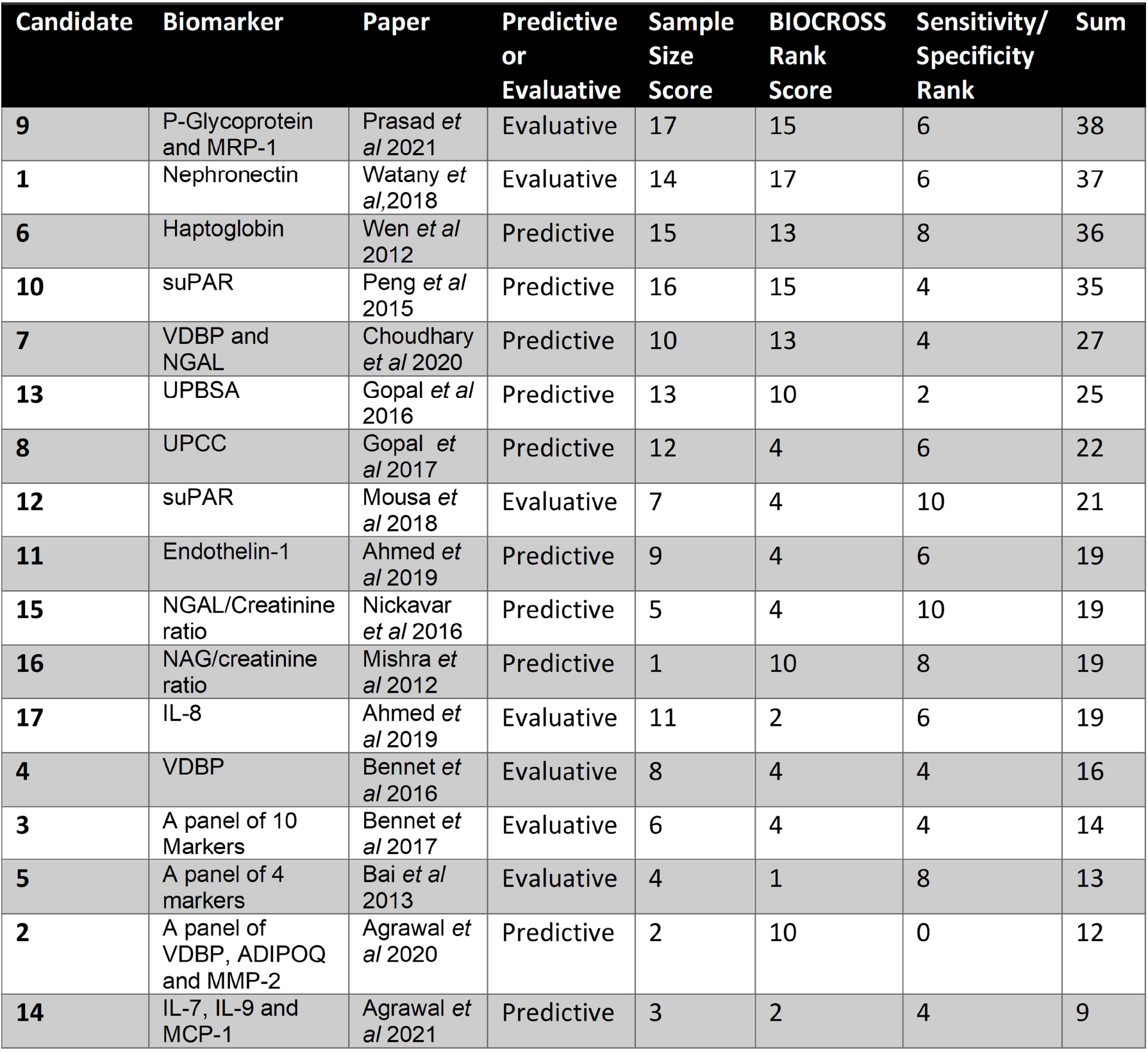
Candidate Biomarker Ranking. The biomarkers and panels of biomarkers identified in the selected manuscripts were scored according to the criterion at the top of each column. They were then ranked to help identify the most promising biomarkers.

The biomarkers were ranked according to their sample size which ranged from 18 participants for a study of the NAG/creatinine ratio to 254 participants for a study on P-glycoprotein and MRP-1. The biomarkers were also ranked by their BIOCROSS score giving the lowest scoring paper (candidate 5 [23]) point and the highest scoring paper (candidate 1 [19]) 17 points. Additionally, biomarkers were scored by their sensitivity/specificity according to the ranges they fell within (Figure 4). Papers that did not disclose sensitivity/specificity values received 0 points.

## Discussion

The field of clinical nephrology is working toward finding predictive biomarkers for SRNS to save patients being exposed to futile steroid treatments. Unfortunately for these kinds of studies INS as a whole is rare and SRNS even more so. The importance of undertaking studies with an adequate sample size is demonstrated by the two studies for candidate biomarkers 12 and 15 [30, 33]. Both reported 100% diagnostic accuracy for the candidate markers under investigation (serum suPAR as an evaluative marker and NGAL/creatinine ratio as a predictive marker). These molecules need now to be evaluated prospectively. It was recently done for suPAR in the prediction of outcomes in septic acute kidney injury [120]

We did not set a lower limit for sample size. Nephrotic syndrome and its subdivisions are rare diseases hence recruitment can be difficult. We accept that smaller sample sizes of patient groups are likely to be less representative and more prone to error however, we wanted to generate a list of candidate biomarkers in the field for further validation.

Overall, studies were considered to have low risk of bias. Three studies were rated as having an unknown risk of bias [20, 23, 32]), which was because of an inadequate description of the exclusion criteria, and as such it is impossible to determine how generalisable their study group is and how well the studies addressed potential confounding factors.

The main limitations of studies reported to date include sampling method, study design, application of cut-off threshold, investigator blinding, and timing of testing.

None of the included studies reported a sampling method for patient recruitment. Given the rarity of INS it is likely that studies used consecutive rather than random sampling to recruit patients, but this is not evaluable. Choice of study design is also a concern. Most identified in this review used a case-control design. It is preferable to avoid case-controlled study design, because it can be difficult to identify an appropriate control group to reduce risk of bias [121]. However, such designs have practical advantage for recruiting cases when the disease is rare. The reporting of control selection is therefore critical in order to assess risk of bias.

None of the selected papers pre-specified the threshold cut-off. By selecting the cut-off after the analysis the data is shown in its best light and is therefore likely to lead to an overestimation of the abilities of the index test. Since all the papers that used a cut-off failed to pre-specify, they remain comparable. However, it is worth pointing out that the same cut-offs would be unlikely to yield the same sensitivity/specificity values in a new cohort. Where possible, authors should pre-specify the cut-offs to increase the accuracy of their sensitivities and specificities.

Future studies are also encouraged to undertake assessor blinding, which enables the impartial analysis of the index test. None of the studies included in this review provided any details about blinding of the results of the index test of reference standard.

It is also important to report when the index test was performed relative to when the samples were taken. Progression of the pathophysiology will have an impact on the abundance of the biomarkers being tested. Again, to make an accurate assessment of risk of bias studies need to be given all the information. Almost half of studies included in this review (8/17) did not report any information regarding when the index test took place relative to the reference standard.

To improve the clinical management of patients with steroid resistant nephrotic syndrome, the goal is to identify predictive markers that will obviate the need to expose these patients to toxic, ineffective treatment.

Candidate biomarkers identified in this review were ranked according to a combination of their sample size, the BIOCROSS score and their sensitivity/specificity. Though P-glycoprotein and MRP-1 scored the most according to these criteria, it is less clinically useful since these are evaluative rather than predictive biomarkers. Haptoglobin was the most rigorously tested predictive marker with the most promising sensitivity/specificity values, closely followed by suPAR.

The biggest flaw with the current evidence-base is how reliant these biomarker studies are on case-controlled studies. Many of the limitations of the studies included here could be overcome by adopting the PRoBE (Prospective-Specimen Collection Retrospective Blinded Evaluation) approach using a biobank such as NURTuRE-INS, which currently contains samples from 742 INS patients [122, 123]. NURTuRE-INS is a well-defined prospective cohort that collects blood and urine from both steroid resistant and steroid sensitive idiopathic nephrotic syndrome patients. The bio samples are collected during periods of active disease relapse) and remission providing vital internal controls for each patient. Despite the multicentre approach, samples are all handled to the same exacting protocol. The 23 renal centres across the UK collect, process, and freeze the samples at -80°C within 2 hours. There is a chronic kidney disease arm of NURTuRE that could provide an additional source of control samples. These could control for markers of inflammatory processes common to multiple kidney diseases.

The PRoBE approach deals with several biases that can be inherent with retrospective case-control study designs. Spectrum bias occurs when case-patients with clear cut examples of the disease (usually severe and/or well-documented) are compared with carefully selected, particularly healthy controls [124]. The manuscripts selected for review here did not describe how patients were sampled (e.g., consecutive, or random), without such information it is difficult to judge the risk of spectrum bias. However, this bias can be avoided when using the PRoBE approach. Subjects in the cohort are identified as patients or controls, then the study group is randomly selected from these sub-groups.

Additionally, by using nested subgroups the discovery and evaluation phases of biomarker identification can be carried out in the same population [123].

suPAR and haptoglobin have emerged from this systematic review as the most promising biomarkers for the prospective distinction between steroid resistant and steroid sensitive variants of idiopathic nephrotic syndrome. It is our strong recommendation that work continues to investigate the utility of these markers using the PRoBE approach on a cohort such as NURTuRE-INS.

## Notes

### Competing Interest Statement

The authors have declared no competing interest.

